# Integration of zebrafish pineal transcriptomes reveals cell type-specific timing

**DOI:** 10.64898/2026.06.07.728976

**Authors:** Yair Wexler, Dengfeng Huang, Jun Yan, Yoav Gothilf

## Abstract

The teleost pineal gland is an eye-like photoreceptive organ with a central role in the circadian clock system, primarily through its melatonin-producing photoreceptor cells. However, the functional molecular interactions between pineal photoreceptors, accessory cells predicted to support photoreceptor function, and projecting neurons remain incompletely understood. Here, we integrated single-cell zebrafish pineal transcriptomes with bulk circadian and light-response pineal transcriptomes. Combined analysis of two single-cell datasets identified novel photoreceptor and neuronal subtypes, including *parietopsin*-expressing cone-like cells and neurons expressing markers of neuronal maturation. Integration with the light-response dataset revealed light inhibition of photoreceptor opsin genes. Integration with circadian transcriptomes from wildtype fish and fish expressing the clock-disrupting dominant-negative CLOCK (ΔCLK) in pineal photoreceptors revealed cell-type-specific rhythmicity. Despite comparable expression of ΔCLK, photoreceptor subtypes differed in sensitivity to rhythm disruption, with rod-like cells (rods) most severely affected. In neurons, despite the absence of ΔCLK expression, rhythm disruption was comparable to that of rods. Moreover, rhythmic neuronal markers and rhythmic photoreceptor markers exhibited a similar circadian pattern, peaking mainly during the early night. These observations suggest that clock function in neurons depend on photoreceptor output. In contrast, accessory cell rhythmic markers were relatively resistant to ΔCLK disruption and peaked predominantly around subjective dawn, consistent with partially autonomous clock function. To facilitate comparative analysis of gene expression, rhythmicity and light responsiveness across pineal cell types, we developed the Zebrafish Pineal Transcriptomics Viewer. Our findings reveal a temporally structured and functionally heterogeneous organization of the zebrafish pineal gland.

## Introduction

In the animal kingdom, timekeeping is governed largely by the circadian clock system. This function relies both on internal synchrony among components of the circadian clock system and on environmental cues for entrainment [1]. The most reliable of these cues is the daily cycle of light and darkness, and timekeeping is therefore closely related to photoreception [2]. Cellular timekeeping is performed by the molecular clock, a set of transcriptional-translational feedback loops that drive rhythmic gene expression and cellular functions in a predictable manner [3], and photoreception is carried out by light-sensitive proteins, such as opsins.

In vertebrates, coupling between photoreception and timekeeping varies between taxa. In mammals, these functions are performed by distinct autonomous structures: the retina detects light, and the hypothalamic suprachiasmatic nucleus (SCN) functions as a central pacemaker, synchronizing peripheral molecular clocks [1]. Although the SCN relies on retinal input for entrainment, it maintains biological rhythms under constant darkness in the absence of retinal input [4,5]. In contrast, other vertebrates exhibit alternative organizational strategies. Teleosts, in which no functional counterpart to the mammalian SCN has been identified, provide a stark example. As exemplified in zebrafish, teleosts possess widespread directly light-entrainable clocks, present in multiple brain regions [6] and across a broad range of peripheral tissues [7,8,9]. Paradoxically, at the center of this seemingly decentralized clock system lies the photoreceptive pineal gland.

The teleost pineal gland is an eye-like photoreceptive organ located at the dorsal surface of the brain beneath the “pineal window”, where reduced pigmentation allows direct light access [10,11]. Phototransduction is mediated by the pineal photoreceptor cells, categorized as cone-like or rod-like based on the expression of opsin and phototransduction genes: cone-like cells ( “pineal cones”) predominantly express the red cone opsins Opn1lw1 and Opn1lw2, whereas rod-like cells (“pineal rods”) express Exorhodopin (Exorh) [12], a homolog of Rhodopsin (Rho), the most studied opsin of the vertebrate retina [13]. As in the retina, pineal photoreceptors are seemingly supported by accessory cells, which share genetic similarities to the retinal pigment epithelium (RPE) and the Müller glia [12]. In additional, projecting neurons innervate brain regions that are also targeted by retinal and cerebellar projections [14,15]. The pineal gland has long been considered a key component of the circadian clock system, as evidenced by the nighttime-restricted production of melatonin [11,14] and by its role in maintaining larval behavioral rhythms under low illumination [16,17]. Despite its established role in the circadian clock system, the functional interactions between distinct cellular populations of the pineal gland, particularly in the context of molecular aspects of circadian timekeeping and light responsiveness, remain poorly defined.

To address this, we combined published and expanded single-cell transcriptomes of the adult zebrafish pineal gland with bulk circadian and light-response pineal transcriptome datasets. Mapping circadian and light-responsive transcriptional dynamics onto these cell types reveals distinct patterns across cellular populations, including previously unrecognized photoreceptor and neuronal subtypes. Based on molecular correlates, we suggest that neuronal circadian clocks likely depend on photoreceptor output, consistent with the role of photoreceptor molecular clocks as a central circadian pacemaker, whereas accessory cell rhythms appear relatively independent and follow a distinct schedule. To facilitate systematic exploration of the integrated datasets, we developed an interactive viewer that enables gene analysis and comparison across all transcriptomes.

Our findings reveal a heterogeneous and temporally structured organization of gene expression in the zebrafish pineal gland. This integrated framework provides new insights into how a single photoreceptive organ mediates both circadian timekeeping and direct photic signaling at the cellular level.

## Materials and Methods

### Zebrafish lines and husbandry

Adult fish of the transgenic line *aanat2^Y9^* (*aanat2:EGFP*) were used in this study to collect samples of the pineal gland for single-cell mRNA sequencing. Adult fish were maintained at 28 °C under 14 h-10 h light/dark (LD) cycles. Husbandry and experimental procedures were approved by the Animal Use Committee of the Institute of Neuroscience, Chinese Academy of Science (permission number NA-045-2019).

### Generation of nighttime single-cell transcriptome

Adult *aanat2^Y9^* fish, in which the pineal gland is marked by eGFP [18], were maintained under 14 h:10 h light/dark (LD) cycles. At zeitgeber time (ZT) 15, 1 h into the dark phase, fish were anaesthetized in 1.5 mM tricaine, sacrificed by decapitation, and their pineal glands were dissected under a fluorescent microscope. Twelve pineal glands were pooled for subsequent processing.

The pooled tissue was dissociated in 500 μl papain solution (20 units/ml) for 20 min at 37℃. Cells were dissociated by gentle pipetting, resuspended, and washed in 1 ml washing solution [19], prepared by adding 650 μl 45% glucose (Invitrogen, USA), 500 μl 1 M HEPES (Sigma, Japan), and 5 ml FBS (Gibco, USA) to 93.85 ml 1x DPBS (Invitrogen, USA). Cells were subsequently washed once in 1x PBS containing 200 mg/ml BSA (NEB, B9000S). filtered through a 20 μm cell strainer, resuspended in PBS/BSA, and counted using a hemocytometer. Cell viability was assessed using trypan blue staining. After confirming >75% viability, cells were loaded onto the 10x Genomics Chromium platform (Single Cell 3’ v3 chemistry) at a concentration of 300 cells/μl.

A total of 5,874 cells were initially obtained and sequenced at a median depth of 20,850 reads per cell, detecting an average of 1,424 genes per cell. Raw sequencing data were processed using the 10x Genomics Cell Ranger pipeline for barcode processing, UMI counting, and generation of filtered gene-barcode matrices. Reads were aligned to a custom Danio rerio GRCz11 reference genome. Removal of low-quality cells and non-pineal cells is described in the following section.

### Transcriptome integration and analysis

#### Combination and analysis of single-cell transcriptomes

Two single-cell mRNA sequencing datasets of adult zebrafish pineal glands were combined to identify novel cell types and their marker genes: previously published data generated from tissues collected during daytime [12] and new data generated from tissues collected at night. Analysis was performed in R using Seurat v3 [20]. Cells were screened by number of featured genes (nFeature_RNA), total mRNA count (nCount_RNA) and percentage of mitochondrial gene (percent.mt). Cells with nFeature_RNA between 350 and 6500, nCount_RNA under 22000 and percent.mt under 0.25 were kept. These values were selected for both datasets. The datasets were then log-normalized.

To identify the pineal cell types, gene expression in the two datasets was aligned using a small subset of “anchors”, pairs of cells from each dataset contained within each other’s neighborhood [20], followed by dimension reduction using Canonical Correlation Analysis (CCA) [21]. This method led to the identification of the 10 cell types listed in the Results. A few contaminating pigment cells, epidermal cells and habenular cells were also identified and filtered out.

Following clustering, the unified dataset was separated back into the day and night datasets, and the process of identifying cell-specific marker genes was carried out for each dataset separately (see Discussion). A gene was defined as a cluster marker if its expression in that cluster was significantly higher than in all other cell types, using Model-based Analysis of Single Cell Transcriptomics (MAST) [22]. Due to the separation of the datasets, the expression of certain cluster markers could only be detected during the day or during the night.

#### Reanalyzing bulk circadian and light-response transcriptomes

The bulk circadian transcriptome datasets were generated from pineal glands of adult fish (wildtype, WT) [23] and Tg(*aanat2*:eGFP-ΔCLK) (“ΔCLK”) [16] fish, that were collected under constant darkness every 4 hours across 48 hours. We set to reanalyze these published datasets and expand upon them. The analysis goals were identifying rhythmic genes in WT pineals, identifying rhythm disruption in ΔCLK, and determining the temporal phase distribution of the pineal gland.

To address the low temporal resolution for detecting gene rhythmicity the cosinor approach was applied to the WT data [24]. For each gene we used ordinary least squares to fit a sine and cosine curve to its temporal transcription profile, assuming a fixed 24-hour period. Let *β_1_* be the regression coefficient of the cosine term and *β_2_* of the sine term, the amplitude was estimated by 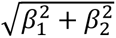, and the phase was estimated by *arctan* (− *β*_2_/*β*_1_). To detect rhythmicity, we tested the amplitude against the Rayleigh distribution, with the scale parameter estimated by the squared root of the mean squared error (MSE). Rhythm disruption in ΔCLK in the subset of identified rhythmic genes was determined by testing for reduced g-factor in ΔCLK compared to WT, using a designated permutations test. G-factor is the ratio of the squared amplitude of the frequency corresponding to 24 hours to the sum of powers of all frequencies following a fast Fourier transform of the signal [25]. All p-values were adjusted using the BH procedure to control the false discovery rate using a hierarchical two step method [26,27] - first the rhythmicity test p-values and subsequently the disruption test p-values.

Determining the temporal distribution of phases was done by hierarchical clustering of the phases of all detected rhythmic genes, converted from radians to sine and cosine terms. An initial 12-cluster distribution was generated and subsequently consolidated into 4 clusters, as shown in **Figure 4**. The full 12-cluster distribution is provided in the supplementary material, Figure S2.

The bulk light-response dataset was generated from pineal glands of adult WT zebrafish placed under constant darkness and then exposed to a 1 h light pulse [28]. Comparing gene expression between the exposed pineals and unexposed control pineals was performed in R using DESeq2 [29,30]. P-values were adjusted using the BH procedure [26], with values under 0.1 considered significant.

#### Comparing cell-specific outcomes

Comparing measures extracted from bulk mRNA datasets between cell types identified by single-cell analysis was based on merging lists of markers. For example, a gene identified as rhythmic in the circadian transcriptome and as a cell-specific marker was defined as a rhythmic marker of those cells. The cell-specific phase distribution was determined from this merged list of rhythmic markers. Comparing marker rhythmicity, rhythm disruption among rhythmic genes, light-response, light-induction among responsive genes and phase distribution between cells was performed using Chi-squared tests. Genes identified as markers of cell types in more than one functional group were removed from the analysis to reduce bias due to overlapping clusters (see Discussion). Comparisons were performed between functional groups (e.g, photoreceptors vs accessory cells) and within each group (e.g. rod-like vs Opn1-expressing cone-like withing photoreceptors). All p-values were adjusted by the BH procedure [26] to control false discovery rate at the 0.05 level.

## Results

### Combining zebrafish pineal single-cell transcriptomes reveal new cell types

Previously published single-cell mRNA sequencing data from the zebrafish pineal gland were generated from tissues collected during the daytime [12]. In that study, seven cell types were reported. Here, we combined this dataset with a new dataset generated from adult pineal glands collected at night (Materials and Methods). After filtering, the combined dataset comprised 8560 cells, 56% of which were derived from the new nighttime dataset.

Clustering of the combined dataset identified 10 distinct cell types (Figure 1), grouped into four functional categories: photoreceptors cells (∼62% of cells), accessory cells (22%), projecting neurons (2%) and non-parenchymal cells (14%). We focus here on the first three groups.

**Figure 1.**
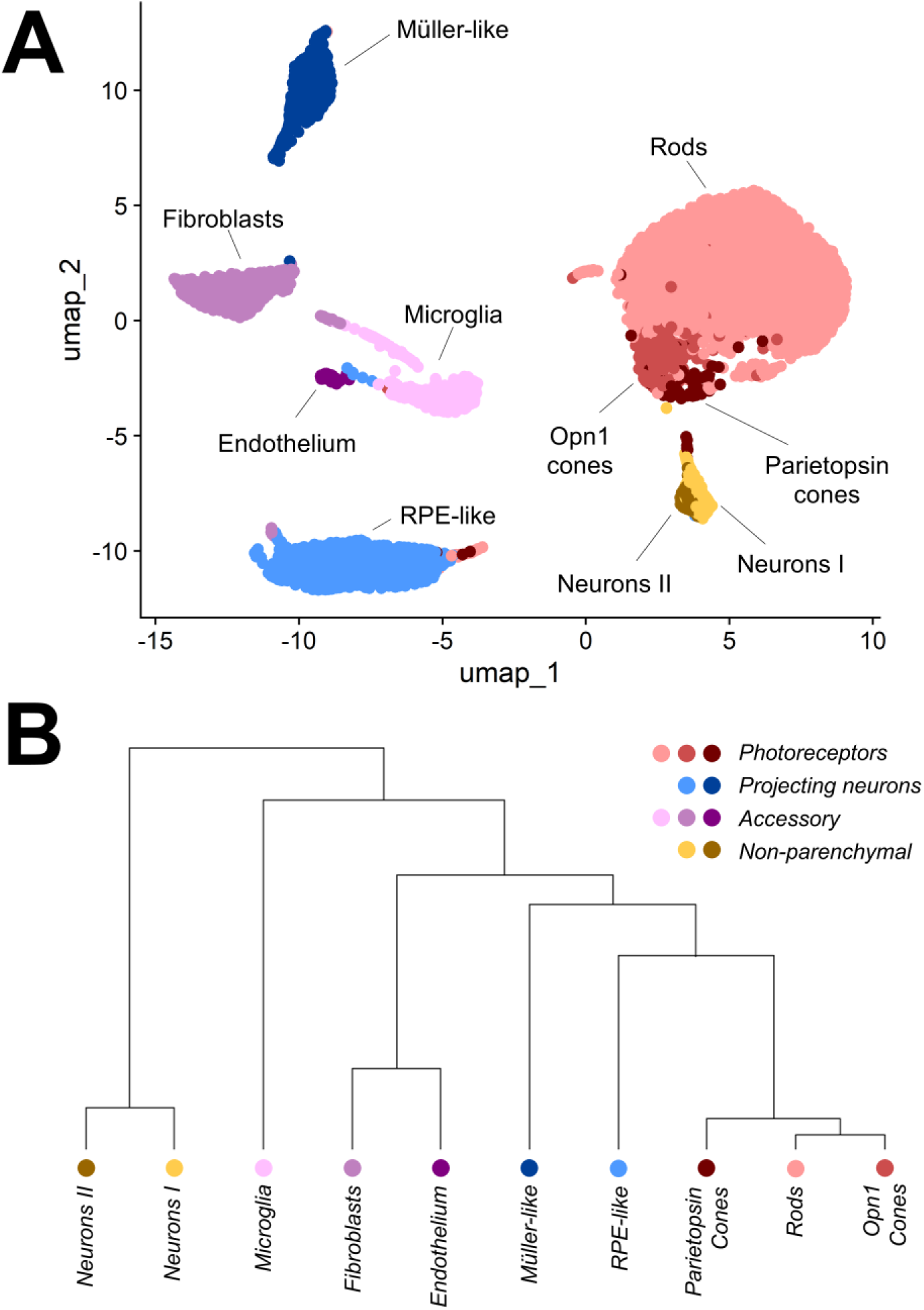
Single-cell mRNA sequencing analysis of the zebrafish pineal gland reveals 10 cell types. A) Visualization of cellular similarities using UMAP. The 10 cell types were groups by predicted functionality: (i) ***photoreceptors*** rods (light red), Opn1 cones (red) and Parietopsin cones (dark red), (ii) ***projecting neurons*** I (orange) and II (brown), (iii) ***accessory cells*** RPE-like (light blue) and Müller-like (dark blue), and (iv) ***non-parenchymal cells*** microglia (pink), fibroblasts (purple) and endothelium (dark purple). B) Although grouped by predicted functionality, a phylogenetic tree of the pineal cell types reveals strong similarities in expression within the photoreceptor group and within the projecting neurons group. Weaker similarity was observed for the accessory subtypes RPE-like and Müller-like cells and within the non-parenchymal subtypes.

Most cell types correspond to those previously described [12]. We nevertheless summarize their identification in this section and extend it in order to provide context for subsequent analysis of cell-specific light response and rhythmicity. We further describe a distinction between two cone-like subtypes and a novel projecting neuron cluster. Non-parenchymal cell types (microglia, fibroblasts and endothelial cells) are described in the supplementary material. The list of cell type-specific markers is provided in Supplementary Table S1.

#### Photoreceptors

Three separate photoreceptor subtypes were identified: (i) rod-like, (ii) Opn1-expressing cone-like, and (iii) Parietopsin-expressing cone-like. Photoreceptors broadly expressed opsins and the melatonin synthesis enzymes *tph1a*, *ddc*, *aanat2*, and *asmt*. For brevity, rod-like and cone-like cells are referred to hereafter as rods and cones, respectively.

As previously described [12], the pineal rods and cones were distinguished primarily by the expression of specific opsins and transducin subunits. Rods expressed the pineal-specific rhodopsin homolog exorh [31,32] and rod-associated Transducin subunits (*gnat1*, *gnb1a/b*, *gngt1*) [12] and constituted the predominant population (58% of cells). Cones expressed cone-specific Transducin subunits (*gnat2*, *gnb3a/b*, *gngt2a*) [12]. The combined analysis distinguished two cone subtypes previously treated as a single population. Opn1-expressing cones (3% of cells) predominantly expressed red cone opsins (*opn1lw1/2*). In contrast, Parietopsin-expressing cones (1.5%) expressed the opsins *parietopsin* and *parapinopsina*, both implicated in color opponency, a mechanism underlying color discrimination, and are thought to be involved in dawn/dusk sensing [33,34,35]. These cells also expressed established cone markers (e.g., *guca1c*, *rx1*, *ndrg1b*), supporting their classification as a distinct cone-like subtype. However, they have a mixed molecular signature, as their arrestin and recoverin profiles showed partial overlap with rod markers (*sagb* and *rcvrna*).

#### Projecting neurons

Projecting neurons comprised two populations, Neurons I (1.7% of cells) and Neurons II (0.85%), both expressing canonical neuronal markers (e.g., *elavl3*, *elavl4*) and the neuroendocrine marker *chga* [36]. Neurons II represent a novel, small cluster that was not detected in the daytime dataset. This population was characterized by expression of neuronal differentiation and axonal elongation factors (e.g., *neurod2*, *neurod6a/b*, *bhlhe22*) [37,38], suggesting a differentiating neuronal state. In addition, Neurons II also co-expressed multiple GABA and glutamate receptor subunits, possibly reflecting responsiveness to both inhibitory and excitatory inputs.

#### Accessory cells

Accessory cells comprised two previously described populations: retinal pigment epithelium (RPE)-like (14% of cells) and Müller-like (8%) [12]. RPE-like cells were characterized by expression of retinoid cycle genes (e.g., *lrata*, *rpe65*, *rdh5* and *cyp27c1*), along with *rlbp1* and the RPE-associated opsins *rgra* and *rgrb*, consistent with a role in retinoid processing [39,40,41]. These cells additionally expressed the opsins *rrh* and *tmtops2b* and were the only non-photoreceptor cell type to exhibit opsin-marker expression. Müller-like cells expressed markers associated with retinal Müller glia, including *vim*, *dkk3b*, *clu*, and *fabp7b*, all associated in vertebrates with retinal response to stress and injury [42–45].

### Cell type-specific transcriptomic response to light corresponds to opsin expression

Following single-cell clustering and identification of pineal cell-type-specific markers, we assessed cell-specific light responsiveness, either induction or inhibition, by integrating the single-cell transcriptomes with a bulk light-response transcriptome [28], generated from pineal glands of dark-entrained wildtype (WT) fish exposed to a 1 h light pulse (Materials and Methods). Light responsiveness was defined as a significant change in log fold expression between light-exposed and unexposed control pineals. The reanalysis revealed 166 genes that significantly responded to the light pulse, of which 97 were induced (58%) and the remainder inhibited (42%). Cell-specific light-responsiveness was quantified by the proportion of marker genes responsive to light and, among these, the proportion that were light-induced. **Figure 2**A is a volcano plot, showing functional group-specific light-responsive markers. The list of light-responsive genes and their corresponding cell types is provided in Supplementary Table S1. Full statistics are provided in Supplementary Table S2.

**Figure 2.**
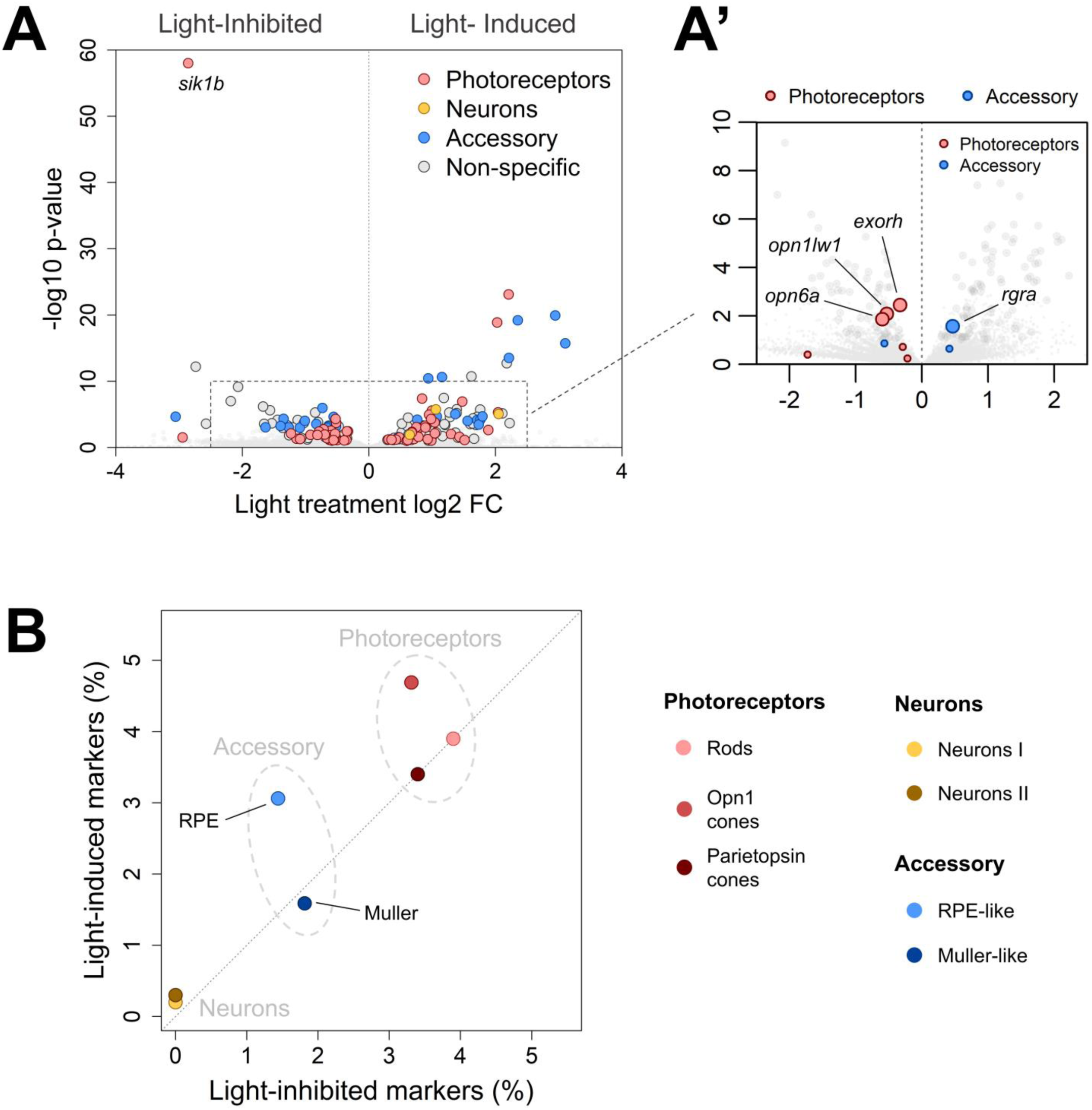
Pineal light responsiveness was primarily observed in photoreceptors, yet despite the predominance of light-induced markers, photoreceptor opsins were consistently light-inhibited. A) Volcano plot showing the effect of the 1 h light pulse on gene expression. The x-axis indicates log_2_ fold-change relative to dark controls, where positive and negative values represent light induction and inhibition, respectively. The y-axis shows - log_10_(p-value), where values above 1 correspond to p < 0.1. Functional group-specific markers are colored pink (photoreceptors), orange (neurons) or blue (accessory cells), and non-specific genes are colored grey. Small grey dots in the background represent non-significant genes. A’) Enlarged view of the volcano plot, corresponding to the dashed square in panel A, highlighting opsin expression. Dot size indicates statistical significance (large: p < 0.1; small: p > 0.1). All six photoreceptor opsins included in the light-response dataset were light-inhibited, including the three significantly responsive opsins *exorh*, *opn1lw1* and *opn6a*, despite overall tendency towards light induction. In contrast, the RPE-like opsin *rgra* was light-induced. B) Proportion of light-induced markers out of all cell type-specific markers (y-axis) relative to the equivalent proportion of light-inhibited markers (x-axis). Values above and below the grey identity line indicate relative enrichment for light-induced and light-inhibited markers, respectively. Photoreceptor subtypes exhibited the highest proportion of light-responsive genes (p = 0.0099** vs accessory; p < 0.001*** vs neurons), whereas neuronal markers were almost entirely non-responsive (p < 0.001*** vs accessory). No differences were detected between photoreceptor and accessory subtypes in the direction of response (p > 0.05). **% light-responsive markers (x: % light-inhibited (↓), y: % light-induced (↑))**: *Rods* 7.8% (**↓** 3.9%, **↑** 3.9%), *Opn1 cones* 8.0% (**↓** 3.3%, **↑** 4.7%), *Parietopsin cones* 6.8% (**↓** 3.4%, **↑** 3.4%), *Neurons I* 0.2% (**↓** 0%, **↑** 0.2%), *Neurons II* 0.3% (**↓** 0%, **↑** 0.3%), *RPE-like* 4.5% (**↓** 1.4%, **↑** 3.1%), *Muller-like* 3.4% (**↓** 1.8%, **↑** 1.6%). P-values were obtained using Chi-squared tests with BH correction (see Materials and Methods). Asterisks represent p-values, where “*” is under 0.05, “**” is under 0.01 and “***” is under 0.001. Non-parenchymal cells were no included in this analysis.

As shown in **Figure 2**B, light-responsiveness across cell types broadly corresponded to opsin diversity. Neuronal markers did not include opsins and were almost entirely non-responsive to the light pulse (0.2-0.3% of all neuronal markers; p < 0.001 compared to other functional groups). Only three neuron-specific markers were light-responsive, and all of them were induced. In contrast, photoreceptors exhibited the highest proportion of light-responsive marker genes (6.7-8.0% of overall photoreceptor markers), significantly exceeding that of the accessory cells (3.4-4.5%; p = 0.0099). Light-responsive markers included the photoreceptor opsins *exorh* (rods), *opn1lw1* (Opn1-expressing cones), and *opn6a* (pan-photoreceptor), as well as the RPE-like opsin *rgra* (**Figure 2**A’). The relative proportions of light-induced and light-inhibited markers among photoreceptor and accessory subtypes were similar to the overall distribution of responsive genes (58% induced, 42% inhibited; p > 0.05).

Notably, despite the overall tendency toward light induction all three significantly responsive photoreceptor opsins were light inhibited (**Figure 2**A’). Negative fold-change, corresponding to light-inhibition, was also observed for the additional photoreceptor opsins included in the light-response dataset, namely *parapinopsinb*, *opn1lw2* and *opn1sw2*, although these effects were not statistically significant. Nevertheless, these observations suggest that opsin expression is consistently suppressed by light in the photoreceptors. In contrast, the RPE-like opsin *rgra* was light induced.

Together, these findings suggest that acute light responsiveness in the pineal gland is mainly a feature of opsin-expressing cell types, whereas neurons exhibit little evidence of direct photic responsiveness.

### Cell type-specific gene rhythmicity

We assessed cell-specific circadian rhythmicity by integrating the pineal single-cell transcriptomes with two published bulk pineal circadian mRNA-sequencing datasets of wildtype (WT) fish [23] and Tg(*aanat2*:eGFP-ΔCLK) (“ΔCLK”) fish, in which pineal photoreceptor circadian clocks are disrupted [16]. These circadian datasets were generated from pineal glands collected every 4 h over 48 h (**Figure 3**A and Materials and Methods). The list of rhythmic markers and corresponding cell types is provided in Supplementary Table S1. Full statistics are provided in Supplementary Table S2.

**Figure 3.**
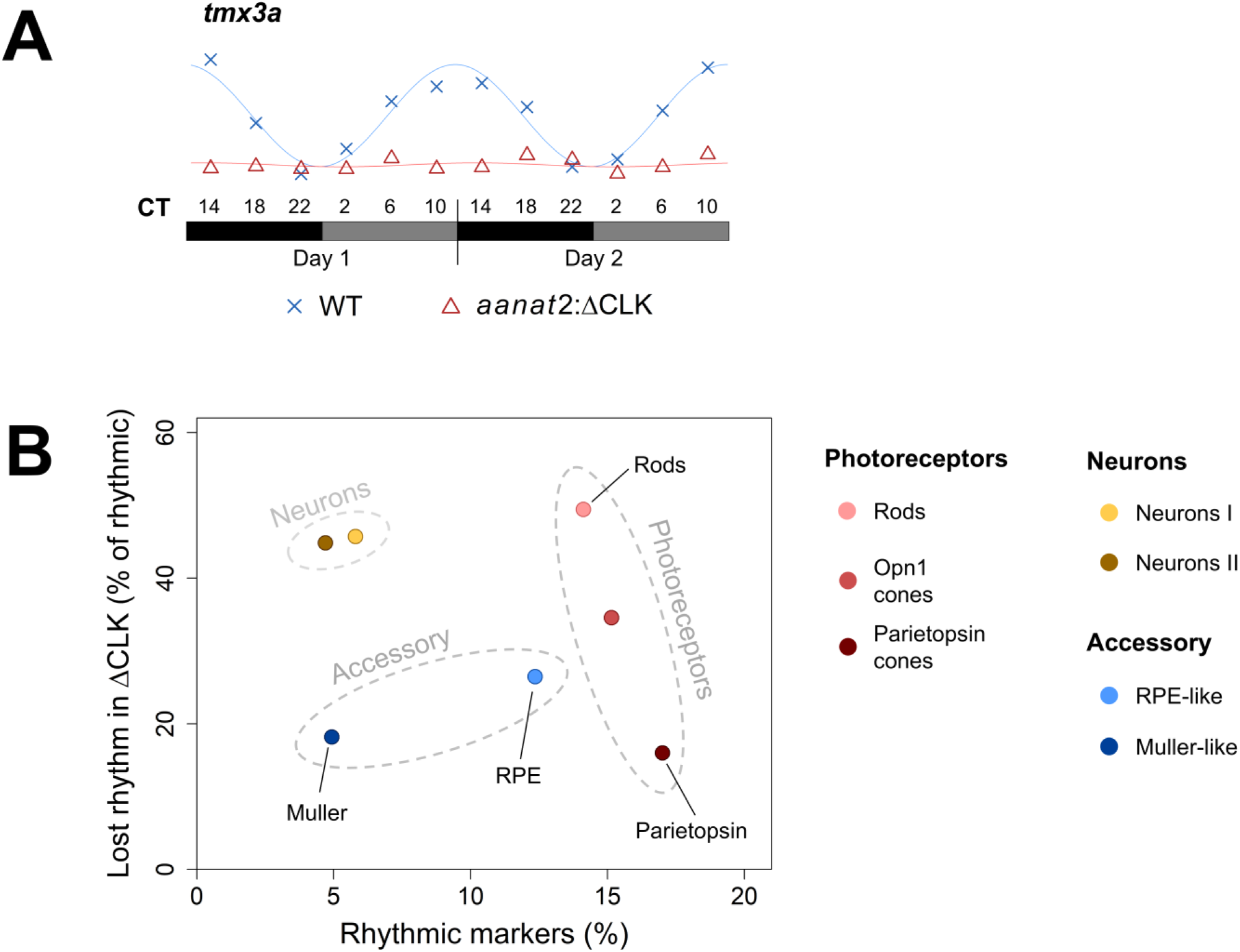
Gene rhythmicity and ΔCLK mutation effect vary between cell types. A) The experimental design and analysis of WT and ΔCLK circadian transcriptomes, illustrated by the expression of *tmx3a*, a rhythmic rod marker whose rhythm was disrupted in Tg(*aanat2*:eGFP-ΔCLK) fish. Pineal glands of adult fish maintained under constant darkness were collected every 4 hours across 48 hours. Black boxes represent subjective night and grey subjective day. Blue crosses mark *tmx3a* expression in WT fish and red triangles expression in ΔCLK fish. The blue and red lines mark the fitted cosinor models (Materials and Methods). *tmx3a* mRNA rhythmicity was completely abolished in ΔCLK fish. B) The x-axis indicates the proportion of rhythmic markers out of all cell type-specific markers in WT pineal glands. The y-axis indicates the proportion, out of those rhythmic markers, of genes whose rhythms were disrupted in ΔCLK fish. Photoreceptors consistently exhibited the highest proportion of rhythmic markers among cell types, but photoreceptor subtypes differed significantly in their sensitivity to ΔCLK disruption (p = 0.021*). Cone markers, particularly those of Parietopsin cones, were less affected by ΔCLK disruption than rod markers (p = 0.010 **, Rods vs Parietopsin cones). Projecting neurons had a relatively low proportion of rhythmic markers but these were strongly affected by ΔCLK, comparable to rods. Accessory cells were heterogenous in rhythmicity (p < 0.001 ***), but both subtypes were relatively resistant to ΔCLK disruption. **% rhythmic markers (% loss of rhythm under ΔCLK)**: *Rods* 14.1% (49.4%), *Opn1 cones* 15.2% (34.5%), *Parietopsin cones* 17.0% (16.0%), *Neurons I* 5.8% (45.7%), *Neurons II* 4.7% (44.8%), *RPE-like* 12.4% (26.5%), *Muller-like* 4.9% (18.2%). P-values for comparison between and within groups were obtained using Chi-squared test and corrected using the BH procedure (see Materials and Methods). Asterisks represent p-values, where “*” is under 0.05, “**” is under 0.01 and “***” is under 0.001. Non-parenchymal cells were no included in this analysis.

Reanalysis of the WT circadian transcriptome (see Materials and Methods) identified 1158 rhythmic genes, 444 of which corresponded to markers of at least one cell type. Cell-specific rhythmicity was quantified by the proportion of rhythmic markers and the effect of ΔCLK expression (**Figure 3**B). Pineal photoreceptors exhibited the highest proportion of rhythmic markers (13% of all photoreceptor-specific markers), significantly exceeding accessory cells (9% of all markers, p = 0.014) and projecting neurons (5% of all neuronal markers, p < 0.001). Accessory cells were heterogeneous (p < 0.001), with RPE-like cells showing higher proportion of rhythmic markers (12%) than Müller-like cells (5%).

In ΔCLK fish, rhythmicity was disrupted in 43% of WT rhythmic genes, with the strongest effect in rods (49% of markers), consistent with the photoreceptor-specific expression of ΔCLK. Cone subtypes were less affected by ΔCLK, particularly Parietopsin cones (16% loss; p = 0.010 compared to rods), despite high expression of *aanat2* and the ΔCLK effector (supplementary material, Figure S1). In contrast, projecting neurons showed substantial disruption (48%) despite lacking ΔCLK expression, suggesting that their molecular clocks may depend on input from the rods. Accessory cells were less affected (27% of rhythmic markers; p = 0.047 compared to neurons), with no differences among accessory subtypes.

These findings suggest that the core clock machinery plays a central role in circadian regulation in pineal rods but a lesser role in cones, and that input from the rods may regulate the molecular clocks of neurons, while accessory cells appear less dependent on photoreceptor clock function.

### Rhythmic photoreceptor and accessory markers peak at different times of day

To assess the temporal organization of pineal functions, we estimated, based on the WT samples, the phases of the 1158 rhythmic genes, defined as the circadian time (CT) of peak expression (see Materials and Methods). Phase clustering identified 4 temporal clusters in sequential order (**Figure 4**A): (I) “Early night”, CT11:23 to CT17:42, (II) “Mid-night”, CT17:42 to CT21:19, (III) “Late night, early day”, CT21:19 to CT4:01, and (IV) “Late day”, CT4:01 to CT11:23.

**Figure 4.**
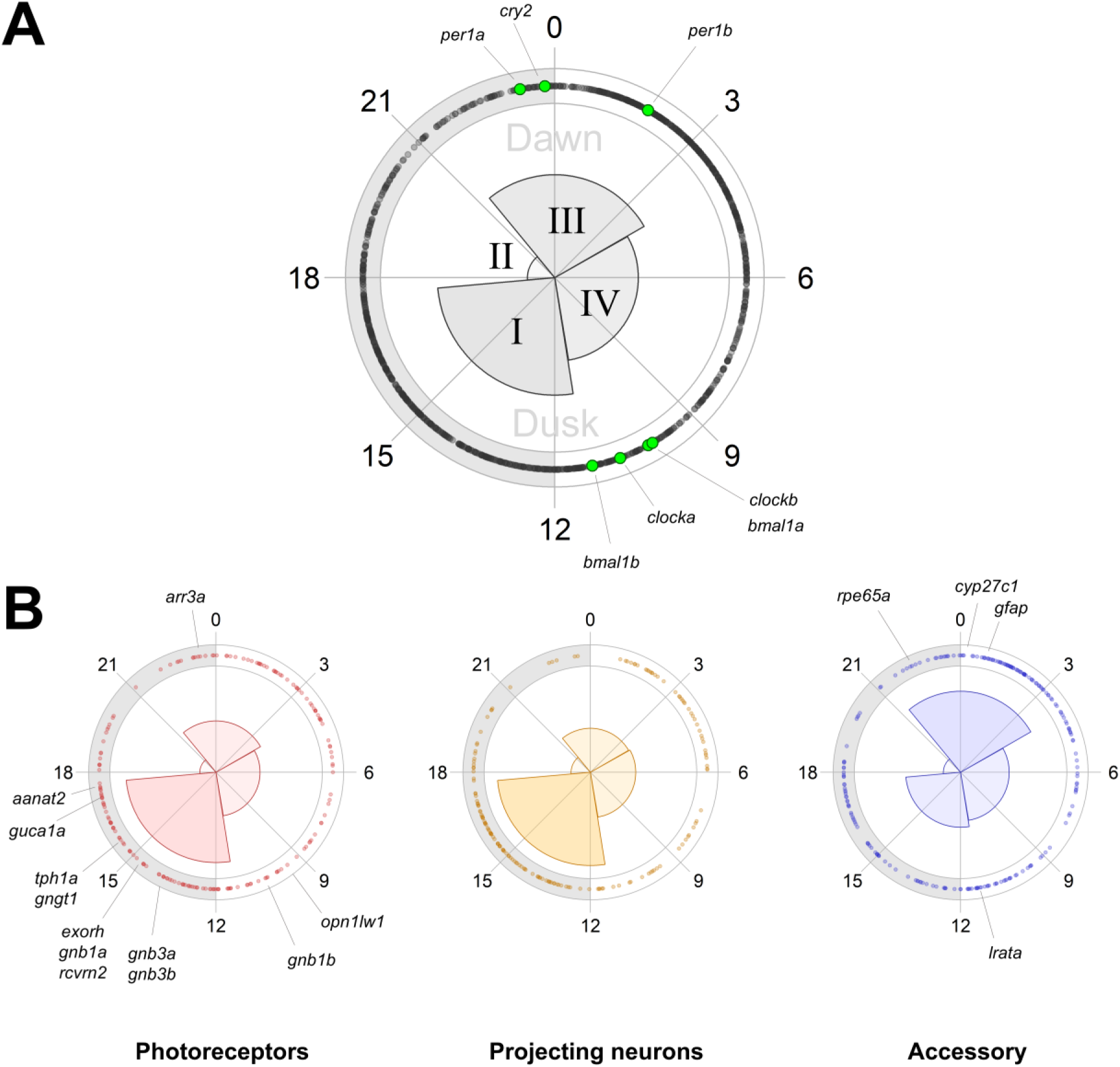
Timing of rhythmic markers varies between cell types. A) Phase distribution of all rhythmic genes identified in the WT circadian transcriptome, divided into 4 temporal clusters: (I) early night, (II) mid-night, (III) late night to early day, and (IV) late day. Slice size represents the proportion of rhythmic genes peaking within each temporal cluster. CT0-12 mark the subjective day (white perimeter) and CT12-24 the subjective night (grey perimeter). Perimeter dots represent individual gene peaks. Cluster I was the largest (35.5% of genes) and II the smallest (8.3%). Core clock genes (green dots) peaked before the subjective dusk (cluster IV) or around the subjective dawn (cluster III). B) Phase distribution for photoreceptors (red, left), projecting neurons (orange, middle) and accessory cells (blue, right). Photoreceptor and neuronal rhythmic markers tended to peak during the early night (I), with 44.7% and 46.2% respectively, whereas accessory markers tended to peak during the late night to early day (cluster III; 40.2%). The accessory temporal distribution differed significantly from that of the photoreceptors (p < 0.001***) and the projecting neurons (p < 0.001***). No difference in distribution was detected between photoreceptors and neurons (p = 0.23) or between subtypes within functional groups (p > 0.05). **Temporal distributions (I, II, III, IV)**: *photoreceptors* (44.7%, 8.0%, 25.4%, 21.8%), *neurons* (46.2%, 9.6%, 21.8%, 22.5%), *accessory* (27.2%, 8.6%, 40.2%, 22.5%). P-values were obtained using Chi-squared test. Asterisks represent p-values, where “*” is under 0.05, “**” is under 0.01 and “***” is under 0.001.

Core-clock genes were primarily distributed between clusters III and IV, with *per1a/b* and *cry2* peaking around dawn (CT23-CT2; cluster III), and *clocka/b* and *bmal1a/b* peaking during the late day, close to dusk (CT10-CT11; cluster IV).

As shown in **Figure 4**B, photoreceptor and projecting neuron markers exhibited similar enrichment in cluster 1, with ∼45% of rhythmic markers peaking during the early night (cluster I). These included genes associated with phototransduction and melatonin synthesis (e.g., *exorh*, *gngt1*, *gnb1a*, *guca1a*, *aanat2*, *tph1a*). Exceptions were the late day (cluster IV) peaks of the cone opsin *opn1lw1* and the rod Transducin subunit *gnb1b,* and the late night (cluster III) peak of the cone arrestin *arr3a*. Accessory cell markers showed a distinct phase distribution (p < 0.001 compared to photoreceptors and neurons), peaking predominantly in the late night and early day (cluster III). These included RPE-like genes implicated in retinal recycling (e.g., *rpe65a*, *cyp27c1*), while *lrata* peaked later in the subjective day. Within each functional group, phase distribution was found similar among cell subtypes (p > 0.05). The list of rhythmic markers, phases and corresponding cell types is provided in Supplementary Table S1. Full statistics are provided in Supplementary Table S2.

Together, these results, along with the effects of ΔCLK support the notion that rod cells synchronize the neuronal molecular clocks, while the accessory clocks maintain, at least in part, an autonomous rhythm. These differential rhythms may reflect distinct clock-driven functions carried by the pineal gland at different times of day.

### Zebrafish Pineal Transcriptomics app identifies *cyp27c1* as a candidate modulator of photoreception

To enable systematic, gene-centered exploration, we developed an interactive viewer that consolidates all transcriptomic measurements into a unified framework. This viewer includes analyzed transcriptomes from several different experiments: the daytime and nighttime wildtype (WT) pineal single-cell transcriptomes, the WT and ΔCLK circadian transcriptomes, a WT light-response pineal transcriptome and a WT light-response brain transcriptome (unpublished).

The viewer enables users to query individual genes and visualize their expression profiles across all included datasets, to compare single-cell distribution, circadian rhythmicity and light-responsiveness between two genes and to generate lists of genes by defined criteria. These criteria include gene name or a part of it, cell-specificity, rhythmicity, impaired rhythmicity in ΔCLK, pineal or brain light-response.

Figure 4 presents an example output from the Zebrafish Pineal Transcriptomics viewer, comparing the expression of *rpe65a* and *cyp27c1* across multiple datasets. Both genes are expressed in the eyes: RPE65 is known as a retinal pigment epithelium marker and a key enzyme in the retinoid cycle, supporting rhodopsin function by regenerating the chromophore 11-cis-retinal [39]. *Cyp27c1* has been implicated in an alternative retinoid pathway in freshwater vertebrates, producing the chromophore 11-cis-3,4-didehydroretinal (vitamin A2), which induces a red-shift in opsin absorption spectra [46,47]. This shift reportedly helps infrared light detection in zebrafish larvae [48]. Interestingly, *rpe65a* and *cyp27c1* exhibited similar patterns across our pineal datasets. Both were enriched in RPE-like cells (**Figure 4**A), both were significantly induced by light (approximately 2-fold increase following the light pulse; **Figure 4**B), and both displayed strong circadian rhythmicity with peaks around subjective dawn (**Figure 4**C). Furthermore, the rhythms of neither gene were impaired by the ΔCLK mutation (*rpe65a*: p = 0.40; *cyp27c1*: p = 0.36), consistent with the above conclusion that RPE-like cells maintain autonomous circadian rhythms. Given the established role of *cyp27c1* in mediating a red-shift in retinal opsin spectral sensitivity, its coordinated expression with *rpe65a* in RPE-like cells and its temporal alignment with dawn suggests a potential role in modulating pineal spectral sensitivity. In this context, *cyp27c1* may contribute to enhanced detection of infrared light and possibly facilitating dawn sensing.

This observation illustrates how the integrated viewer enables identification of genes whose combined cell-type specificity, rhythmicity, and light responsiveness point to potential functional roles that would be difficult to infer from individual datasets alone.

The Zebrafish Pineal Transcriptomic viewer is available at https://gothilf-zebrafish.shinyapps.io/zebrafish-pineal-transcriptomics/.

## Discussion

The zebrafish pineal gland is a photodetector and a circadian pacemaker [11], which drives circadian rhythms within a clock system whose peripheral components are directly light-entrainable [6,8]. The pineal photoreceptors produce melatonin using an enzymatic pathway highly conserved across vertebrates [49]. Melatonin has been the predominant focus in pineal research, owing to its reliability as a signal of the night [49,50]. However, the pineal gland may function as a photodetector in a broader sense.

In zebrafish larvae, pineal dysfunction results in a reduced behavioral response to sudden dimming [51]. Pineal-dependent behavioral responses to sudden dimming have also been reported in Xenopus tadpoles [52,53] and in larvae of blind Mexican cavefish [54]. The short duration of these behavioral responses is incompatible with melatonin signaling. Indeed, the pineal-mediated larval response in zebrafish requires not only photoreceptors but also the projecting neurons [51]. These neurons innervate premotor and precerebellar nuclei in the pretectum, the ventral thalamus and the tegmentum [15] and may therefore directly influence locomotion. A study monitoring *c-fos* expression in pineal neurons in response to changes in illumination showed that most neurons are activated by darkness [11], with a small population of *opn4xa*-expressing neurons responding to green and blue light [55]. Opsin expression in most pineal neurons, however, appears limited. Thus, the extent to which pineal neuron-mediated processes depend on photoreceptor input, and on which of the photoreceptor subtypes, remains unclear.

This uncertainty extends beyond neurons to other cellular components of the pineal gland. While accessory cells are thought to support photoreceptor function [12,56], their contribution to circadian timekeeping and light-dependent processes remains insufficiently characterized. More broadly, how these cell populations interact to generate pineal function remains unclear. In this study, we aimed to provide insights into these interactions by combining multiple single-cell and bulk transcriptomes of the zebrafish pineal gland.

Integration of a newly generated single-cell dataset with a previous one revealed ten distinct cell populations (**Figure 1**), including a novel distinction between two subtypes of cone-like cells (“cones”). The majority of cones express the red cone opsins *opn1lw1* and *opn1lw2*, whereas a smaller population expresses the non-visual opsin *parietopsin*, which has been implicated in dawn sensing [33]. Unsurprisingly, the abundant expression of opsins corresponded to the high light responsiveness of photoreceptor cell types relative to the other pineal populations (**Figure 2**). Notably, however, photoreceptor opsins themselves were consistently light-inhibited despite the overall predominance of light-induced genes following the light pulse. This coordinated suppression may reflect the proposed specialization of the teleost pineal gland for darkness detection and low-light signaling [57,11]. Photoreceptor subtypes also exhibited a high rate of gene rhythmicity compared to other cell types. However, Parietopsin-expressing cones stood out in that their markers were relatively resistant to loss of rhythmicity in ΔCLK-expressing pineal glands (**Figure 3**). ΔCLK is a dominant-negative CLOCKa mutation that disrupts the molecular clock [15]. These findings suggest a degree of independence in clock function between photoreceptor subtypes, as well as a possible variation in molecular clock architecture.

A further distinction revealed by the integrated single-cell analysis is between two subtypes of projecting neurons, Neurons I and II. Interestingly, *opn4xa* was not detected in any of these neurons, and therefore this distinction does not correspond to the previously described dark-and light-activated populations [55]. Neurons II express several markers associated with neuronal maturation, including *neurod2*, *neurod6a/b,* and *bhlhe22* [37,38]. In contrast to the photoreceptors, neuronal markers exhibited almost no responsiveness to the light pulse. Given the involvement of pineal neurons in the behavioral response to sudden darkness [51], future investigation of transcriptomic responses to darkness in neuronal populations may provide additional insight. Consistent with their limited direct light responsiveness, neuronal clock function appeared to depend on rod-like (“rods”) photoreceptor input. This is supported by the pronounced effect of ΔCLK on neuronal marker rhythmicity, despite neurons not expressing the mutant protein, as well as by the alignment of neuronal and photoreceptor phase distributions (**Figure 4**B). However, this interpretation should be treated with caution. Due to the relatively low rate of rhythmicity in neurons and differences in cell proportions, the apparent similarity between neurons and rods may reflect cluster overlap rather than true functional dependence (see below).

Unlike the neurons, accessory cell clocks appeared to be relatively independent of photoreceptors. RPE-like cells exhibited a high rate of rhythmicity but were resistant to loss of rhythmicity under ΔCLK expression relative to rods. Furthermore, rhythmic accessory markers exhibited a significant shift in timing relative to photoreceptors. Accessory markers tended to peak around the subjective dawn, aligning with the phases of *per1a/b* and *cry2*. Among the rhythmic accessory markers resistant to ΔCLK disruption and peaking around the subjective dawn were *rpe65a* and *cyp27c1* (**Figure 5**), both associated with retinoid metabolism and photoreception support [39,48]. In contrast, photoreceptor markers tended to peak in the early night, following the predusk expression of *clocka/b* and *bmal1a/b*. These included *exorh*, several rod and cone transducin subunits, and *aanat2* and *tph1a*, which encode enzymes involved in melatonin synthesis. RPE-like cells expressed opsins of their own, namely *rgra* and *rgrb*, which may contribute to their partial autonomous circadian clock entrainment. Moreover, *rgra* and several RPE-associated visual cycle genes, including *rpe65a*, *cyp27c1* and *rdh5*, were light induced, in contrast to the coordinated light inhibition observed in photoreceptor opsins. This distinction is consistent with light-dependent activation of retinoid recycling and photoreceptor-support functions during the day.

**Figure 5.**
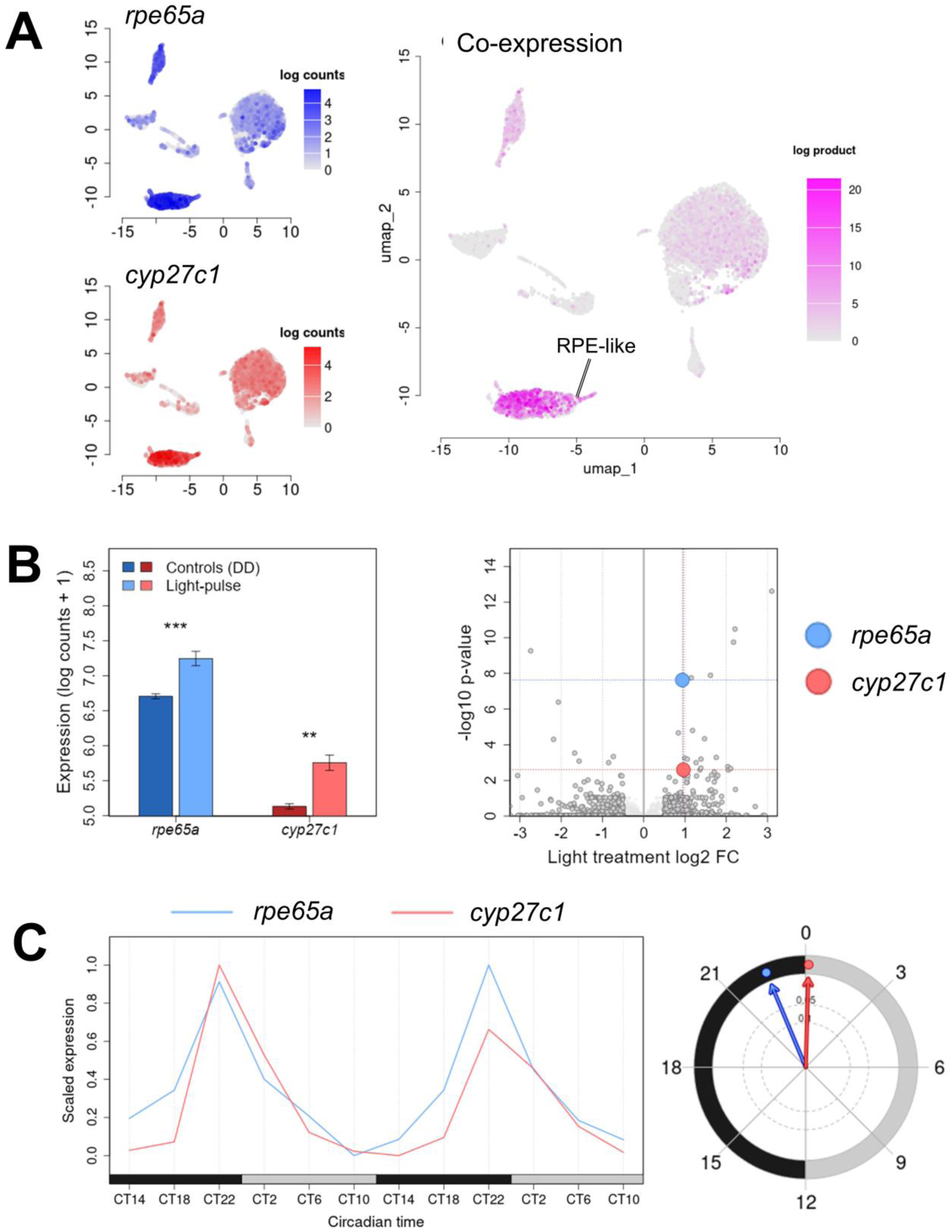
Analysis using the Zebrafish Pineal Transcriptomics viewer reveals similarities between *rpe65a* and *cyp27c1*. A) UMAP plots of *rpe65a* (blue, top left), *cyp27c1* (red, bottom left), and their co-expression (purple, right), with color intensity marking expression level in the single-cell datasets. Both genes were enriched in RPE-like cells. B) Left – bars represent the average expression ± SEM for *rpe65a* (blue) and *cyp27c1* (red) in the light-response dataset. Dark color marks the control group (DD) and bright color the light-pulse group. Right - volcano plot showing the effect of the 1 h light pulse on gene expression. The x axis indicates log_2_ fold-change relative to dark controls, where positive and negative values represent light induction and inhibition, respectively. The y axis shows -log_10_(p-value), where values above 1 correspond to p < 0.1. Grey dots mark all genes that exhibited a log_2_ fold-change of at least 0.5. Both *rpe65a* (blue) and *cyp27c1* (red) were significantly light induced. *rpe65a* exhibited a log_2_ fold-change of 0.94 (log_2_1.91, p < 0.001 ***). *cyp27c1* exhibited a log_2_ fold-change of 0.96 (log_2_1.94, p = 0.003 **). C) Left – the rhythmic expression of *rpe65a* (blue) and *cyp27c1* (red) across 48 hours of measurement under constant darkness in the WT circadian datasets. The grey and black boxes represent the objective day (CT0-12) and night (CT12-0), respectively. Right – the phases of both rhythms. Both *rpe65a* (blue) and *cyp27c1* (red) exhibited robust circadian rhythms (p < 0.001 *** for both), with similar phases around dawn. *rpe65a* peaked at CT22.5 and *cyp27c1* at CT0. Neither gene exhibited loss of rhythmicity in ΔCLK-expressing pineal glands (p > 0.05; not shown).

While further characterization of cell-specific functions is required to define the precise biological processes underlying these phase distributions, the present findings provide strong evidence that photoreceptive and maintenance-related functions are temporally segregated and governed by partially independent, cell-type-specific programs. Thus, the seemingly decentralized nature of the zebrafish circadian clock system may extend to the pineal gland itself, where the photoreceptor population acts as a prominent but not exclusive pacemaker.

Finally, the interactive viewer developed in this study (**Figure 5**) enables systematic exploration of pineal transcriptomic data, allowing direct comparison of cell-type specificity, circadian rhythmicity, and light responsiveness across datasets. This resource facilitates the identification of coordinated gene expression patterns and candidate functional relationships that would be difficult to infer from individual experiments alone.

### Considerations and limitations of integrating transcriptomic data

Integration of single-cell and bulk mRNA transcriptomes has become an increasingly common strategy for identifying cell type-specific characteristics. For example, a similar approach was recently used to predict therapeutic responses based on lymphocyte profiles in gastric cancer [58] and lung squamous cell carcinoma [59]. However, transcriptomic integration is not without limitations. In the present study, datasets were generated from multiple independent experiments that varied in design, fish lines, and sample processing. The inferential capacity of such integrative analyses depends on the degree of ambient mRNA contamination [60], marker specificity, cell distribution, and statistical power. Therefore, the integrated analysis required, in addition to cautious interpretation of the results, careful weighing of different analysis strategies.

On one hand, a stringent approach to cell type identification was necessary as a basis for comparison, to minimize cluster overlap and misclassification, which may disproportionately bias results toward more abundant cell types. Accordingly, as described in the Materials and Methods, we adopted the integration pipeline proposed by Stuart and Satija (2019) [21], aligning the single-cell datasets using a limited subset of shared genes (“anchors”).

On the other hand, maximizing statistical power required a more permissive approach for identifying cellular markers, rhythmicity and light responsiveness. For example, comparisons of cell-specific phase distributions were based on genes identified both as cell-type markers in the single-cell datasets and as rhythmic in the WT circadian dataset. A limited number of genes in either category would substantially reduce statistical power. We therefore prioritized sensitivity, aiming to maximize the number of true positives, even at the cost of increased false positives. This trade-off is partially mitigated by the expectation that genes falsely classified as rhythmic contribute phases that are approximately uniformly distributed throughout the day, thereby introducing noise rather than systematic bias. In contrast, true rhythmic markers are expected to cluster around biologically relevant peak phases. Therefore, despite increased background noise, such a permissive approach improves recovery of the underlying temporal distributions.

For this purpose, anchor-based clustering was followed by marker identification in each single-cell dataset separately using the full gene set (Materials and Methods). This approach recovered many markers that were not detected in the integrated dataset, including known pineal-enhanced genes such as *asmt, ddc* and *tph2*. Consistent with this, previous stringent analysis of the bulk circadian transcriptomes identified 290 rhythmic genes [23], of which 133 were classified here as cell-specific. The present analysis more than tripled this number, identifying 444 cell-specific markers among 1158 rhythmic genes. Likewise, reanalysis of the light-response dataset using DESeq2 doubled the number of identified responsive markers and revealed coordinated light inhibition of photoreceptor opsins. However, in this case the number of responsive genes remained low (166), limiting inference power considerably. Statistical power was further improved by restricting the number of tests, comparing functional groups and cell types within functional groups rather than pairwise comparisons between cell types, as well as excluding non-parenchymal tests.

Nevertheless, the risk of bias towards dominant cell-types, particularly rod-like cells (58% of cells), remains. Projecting neurons, which constitute only 2.5% of cells, showed similarities to rods both in the effect of ΔCLK on rhythmicity and in phase distribution. One interpretation, to be explored in future work, is that rods function as pacemakers for the neuronal circadian clocks. However, neuronal rhythmic markers were scarce and expressed at low levels, in contrast to photoreceptors and accessory cells, which exhibited an abundance of highly expressed rhythmic markers. Thus, overlap between rod and neuronal transcriptional profiles may also account for these similarities. In this context, the evidence for differences in phase distribution between photoreceptors and accessory cells is more robust.

## Data and software availability

The nighttime single-cell dataset was submitted to GEO, accession number GSE333181.

Previously published datasets: daytime single-cell [12], GEO: GSE123778; WT light-response [28], WT circadian [23] and ΔCLK circadian [16], SRA: SRP016132.

## Acknowledgements

We thank Inbal Shainer, Shir Confino, Yotam Elazary and Zohar Ben-Moshe for a fruitful discussion and expert advice, and for their assistance in testing the viewer application.

