## supplementary material for "Integration of zebrafish pineal transcriptomes reveals cell type-specific timing"

Yair Wexler[1,6], Dengfeng Huang[2,3], Jun Yan[2,4] and Yoav Gothilf[1,5]

[1] School of Neurobiology, Biochemistry and Biophysics, The George S. Wise Faculty of Life Sciences, Tel-Aviv University, Tel Aviv, Israel

[2] Institute of Neuroscience, CAS Center for Excellence in Brain Science and Intelligence Technology, Chinese Academy of Sciences, Shanghai 200031, China.

[3] Department of Pathology, Shanghai Tenth People's Hospital Affiliated to Tongji University, Shanghai 200072, China.

[4] School of Future Technology, University of Chinese Academy of Sciences, Beijing 101408, China.

[5] Sagol School of Neuroscience, Tel-Aviv University, Tel Aviv, Israel

[6] Lead contact

### Supplementary results

#### Non-parenchymal cells of the zebrafish pineal gland.

As detailed in the main text, reanalysis and integration of the single-cell datasets expanded upon a previous study in which seven cell types were identified [1]. The integrated analysis identified 10 cells types, grouped into four functional categories: photoreceptor cells (~62% of cells), accessory cells (22%), projecting neurons (2%) and non-parenchymal cells (14%). Non-parenchymal cell types were excluded from the integrated circadian and light-response analyses in order to limit the number of comparisons performed in this study. A brief characterization of these cell types is therefore provided here, together with descriptive statistics but without statistical comparisons to other cell types or functional groups.

The three non-parenchymal cell types identified were microglia (5.5% of cells), fibroblasts (7.5% of cells), and endothelial cells (0.9%). These clusters express canonical markers of their respective cell types. Microglia expressed *cd74a, cd74b, apoeb* and *ctsc*, as previously described [1]. Fibroblasts expressed the pan-fibroblast markers *pdgfra, col1a2* and *dcn* [2,3]. Additional fibroblast markers included *sfrp1a* and *ccl25b*, previously identified in larval reticular cells supporting the vasculature of the caudal vein plexus [2]. Endothelial cells expressed established endothelial markers, including *kdrl, fli1a* and *pecam1* [4]. A small subset exhibited additionally exhibited the parapineal markers *krt4* and *tbx2b* [5], potentially reflecting minor contamination from neighboring parapineal cells.

Non-parenchymal marker genes were the least light-responsive of all cell types (microglia: 1.3%; fibroblasts: 2.9%; endothelium: 2.7%), substantially lower than the next least light-responsive cell types, Neurons I and Müller-like cells (5.6%). Among the light-responsive markers, the proportion of light-induced genes was comparable to that observed in photoreceptors and accessory cells (microglia: 37.5%; fibroblasts: 66.7%; endothelium: 60.0%). Marker rhythmicity in non-parenchymal cells was comparable to that observed in neurons and Müller-like cells, showing overall low rhythmicity (microglia: 5.5%; fibroblasts: 5.9%; endothelium: 6.2%). However, markers of all three non-parenchymal clusters were highly sensitive to ΔCLK disruption (microglia: 47.1%; fibroblasts: 45.8%; endothelium: 43.5%), similar to neurons (44.8%–45.7%) but unlike Müller-like cells (18.2%), suggesting that input from the pineal rods may contribute to regulation of non-parenchymal molecular clocks.

#### Cell-specific expression of *aanat2* and *clocka*

The *aanat2* promoter drives the expression of the dominant negative mutation ΔCLK, a faulty CLOCKa protein, in the pineal glands of Tg(*aanat2*:eGFP-ΔCLK) fish [6]. In **Figure 3** of the main text, we show that 17% of Parietopsin cone markers were identified as rhythmic, the highest proportion among pineal cell types, yet only 16% of these markers lost rhythmicity in ΔCLK fish, the lowest proportion among pineal cell types. Thus, Parietopsin cones stood out both in their strong tendency toward rhythmic expression and in their relative insensitivity to ΔCLK disruption. In contrast, rhythmic rod markers were highly sensitive to ΔCLK disruption, with 49% losing rhythmicity.

The reduced sensitivity of Parietopsin cones may reflect their lower *aanat2* expression compared to rods and Opn1 cones ($p < 0.001$). However, *aanat2* expression remained substantially higher than in any non-photoreceptor cell type (**Figure S1**A), consistent with its role as a pan-photoreceptor marker. Opn1 cones, in which *aanat2* expression was comparable to that of the rods, exhibited intermediate ΔCLK sensitivity (35% rhythm loss), although only the difference between rods and Parietopsin cones was statistically significant. Furthermore, *aanat2* expression in Parietopsin cones exceeded that of *clocka* by orders of magnitude (**Figure S1**B), suggesting that ΔCLK should retain a dominant-negative effect in these cells. Therefore, differences in *aanat2* expression alone are unlikely to fully explain the disparity in ΔCLK sensitivity among photoreceptor subtypes.

Given the high proportion of rhythmic markers in Parietopsin cones, their relative insensitivity to ΔCLK disruption may also reflect a reduced dependence of their molecular clocks on CLOCK-mediated transcription.

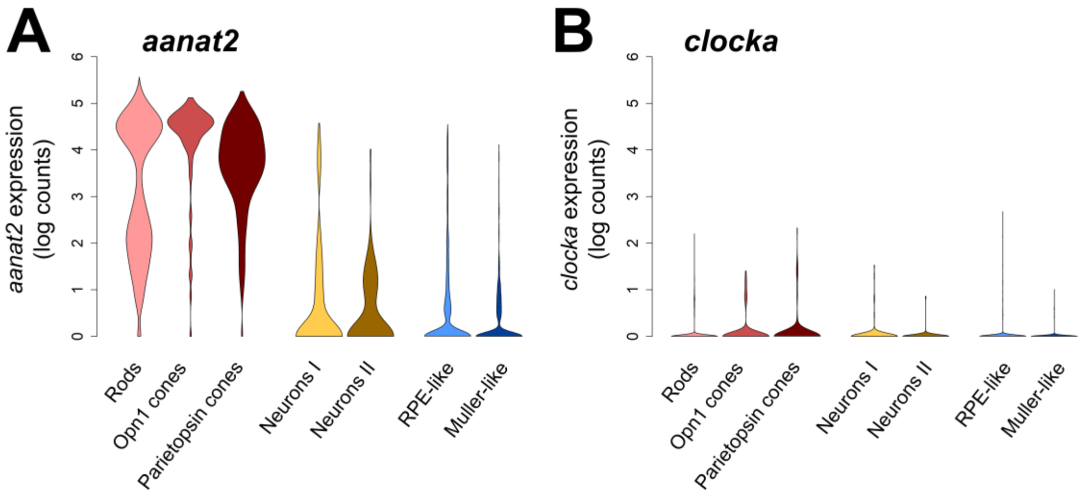


**Figure S1. Supplementary to Figure 3 of the main text: cell-specific expression of *aanat2* and *clocka*.** A-B) Violin plots showing the cell-specific transcriptomic distributions of *aanat2* (left) and *clocka* (right), on a log10 scale. A) *aanat2* expression in Parietopsin cones was generally lower than in rods and Opn1 cones ($p < 0.001$), but substantially higher than in any non-photoreceptor cell type. B) *clocka* expression was relatively uniform across cell types and, in all photoreceptor subtypes, considerably lower than *aanat2* expression, consistent with substantial ΔCLK expression in all photoreceptor populations.

#### 12-cluster phase distribution

The main text presents a comparison between photoreceptors, neurons and accessory cells in the phase distributions of their rhythmic markers. An initial high-resolution 12-cluster temporal distribution (supplementary **Figure S2**) was generated to reflect the temporal resolution of the sampled circadian transcriptome. However, a lower-resolution 4-cluster distribution was ultimately used in the main analysis, as it more clearly captured the major temporal differences between functional groups without excessive subdivision or visual noise. Importantly, the choice of 12 or 4 clusters had little effect on the comparison between functional groups. In both classifications, the predominant temporal cluster in neurons corresponded to that of the photoreceptors, containing rhythmic genes peaking during the early night (CT13:07–15:46). However, this dominant cluster was more concentrated in neurons (26% of rhythmic markers) than in photoreceptors (22%). In contrast, the predominant temporal cluster of accessory cells contained genes peaking during the early day (CT0:16–2:18, 18% of markers), while the overall phase distribution of accessory-cell markers was more dispersed across the circadian cycle than in the other functional groups.

The starting points of the 12 temporal clusters, in circadian time (CT): (*i*) 11:23 (dusk), (*ii*) 13:07, (*iii*) 15:47, (*iv*) 17:42, (*v*) 19:59, (*vi*) 21:19 (dawn), (*vii*) 0:16, (*viii*) 2:19, (*ix*) 4:01, (*x*) 5:23, (*xi*) 7:23, and (*xii*) 8:57. Clusters *i-iii* corresponded to Cluster I ("early night") of the 4-cluster distribution, *iv* and *v* to Cluster II ("mid-night"), *vi-viii* to Cluster III ("late night, early day"), and *ix-xii* to Cluster IV ("late day").

.

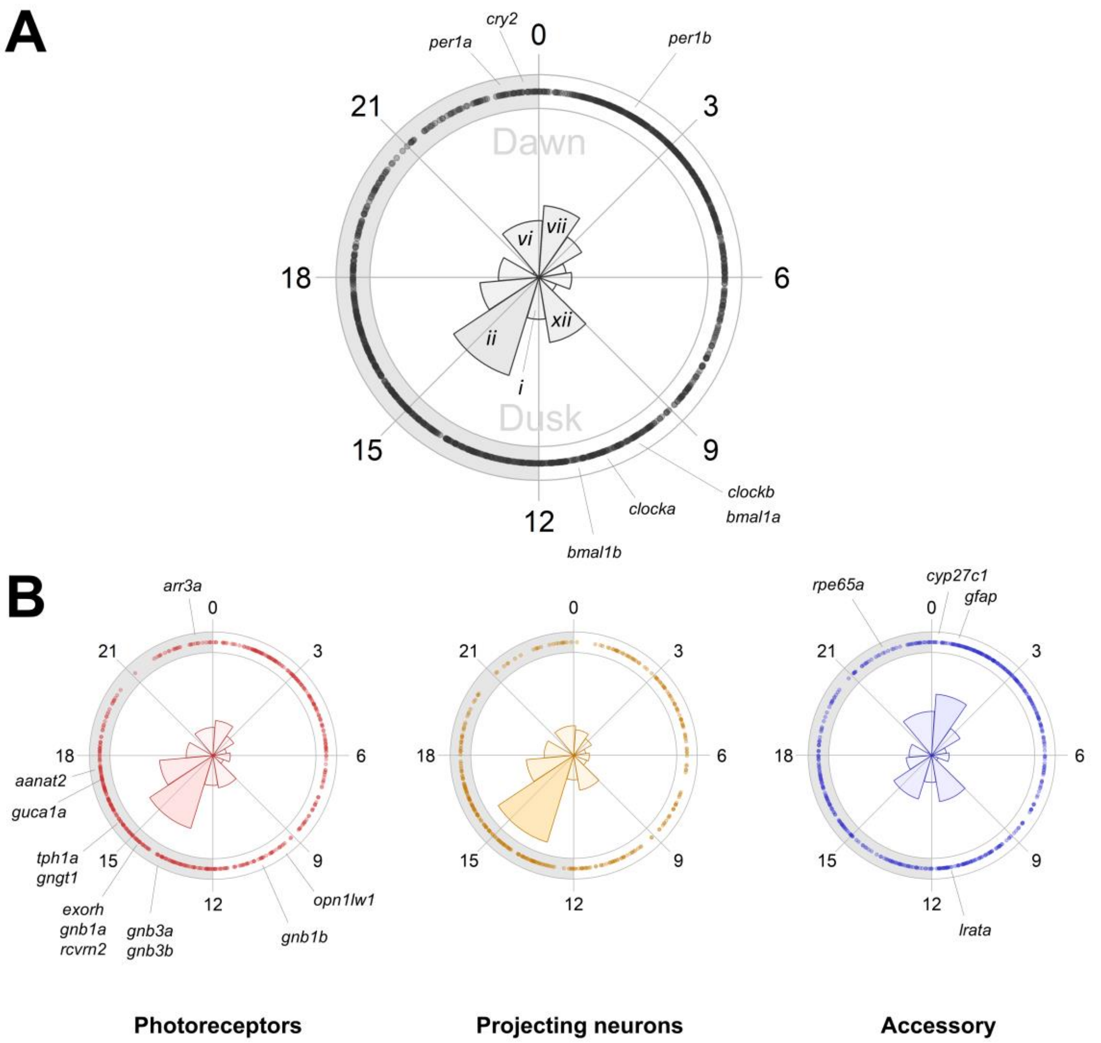


**Figure S2. Supplementary to Figure 4 of the main text: timing of rhythmic markers varies between cell types.** A) The 12-cluster phase distribution of all rhythmic genes identified in the WT circadian transcriptome, beginning at cluster *i* (dusk) and continuing clockwise. Cluster *ii* was the predominant cluster, containing 18% of rhythmic genes. B) Phase distributions of rhythmic markers in photoreceptors (red, left), projecting neurons (orange, middle) and accessory cells (blue, right). Photoreceptor and neuronal markers were enriched in the early night cluster (*ii*), with 22% and 26% of rhythmic markers respectively. Accessory cell markers were enriched in the early day cluster (*vii*; 18%) and showed a broader phase distribution than that the other functional groups.
